# Basset: Learning the regulatory code of the accessible genome with deep convolutional neural networks

**DOI:** 10.1101/028399

**Authors:** David R. Kelley, Jasper Snoek, John Rinn

**Author notes:** Contact information: David R. Kelley, 7 Divinity Ave., Cambridge, MA 02138, Jasper Snoek, School Of Engineering and Applied Science, Harvard University, Cambridge, MA 02138, John L. Rinn, 7 Divinity Ave. Cambridge, MA 02138.

## Abstract

The complex language of eukaryotic gene expression remains incompletely understood. Despite the importance suggested by many noncoding variants statistically associated with human disease, nearly all such variants have unknown mechanism. Here, we address this challenge using an approach based on a recent machine learning advance—deep convolutional neural networks (CNNs). We introduce an open source package Basset (https://github.com/davek44/Basset) to apply CNNs to learn the functional activity of DNA sequences from genomics data. We trained Basset on a compendium of accessible genomic sites mapped in 164 cell types by DNaseI-seq and demonstrate far greater predictive accuracy than previous methods. Basset predictions for the change in accessibility between variant alleles were far greater for GWAS SNPs that are likely to be causal relative to nearby SNPs in linkage disequilibrium with them. With Basset, a researcher can perform a single sequencing assay in their cell type of interest and simultaneously learn that cell’s chromatin accessibility code and annotate every mutation in the genome with its influence on present accessibility and latent potential for accessibility. Thus, Basset offers a powerful computational approach to annotate and interpret the noncoding genome.

## Introduction

The process of identifying genomic sites that show statistical relationships to phenotypes holds great promise for human health and disease (Hindorff et al. 2009). However, our current inability to efficiently interpret noncoding variants impedes progress toward using personal genomes in medicine. Coordinated efforts to survey the noncoding genome have shown that sequences marked by DNA accessibility and certain histone modifications are enriched for variants that are statistically related to phenotypes (Consortium 2012; Ernst et al. 2015). The first stages of a mechanistic hypothesis can now be assigned to variants that directly overlap these annotations (Ritchie et al. 2014; Kircher et al. 2014; Fu et al. 2014).

However, simply considering the overlap of a variant with annotations underutilizes these data; more can be extracted by understanding the DNA-protein interactions as a function of the underlying sequence. Proteins that recognize specific signals in the DNA influence its accessibility and histone modifications (Voss and Hager 2014). Given training data, models parameterized by machine learning can effectively predict protein binding, DNA accessibility, histone modifications, and DNA methylation from the sequence (Benveniste et al. 2014; Lee et al. 2015; Pinello et al. 2014; Das et al. 2006; Setty and Leslie 2015; Whitaker et al. 2015; Arnold et al. 2013). A trained model can then annotate the influence of every nucleotide (and variant) on these regulatory attributes. This upgrades previous approaches in two ways. First, variants can be studied at a finer resolution; researchers can prioritize variants predicted to drive the regulatory activity and devalue those predicted to be irrelevant bystanders. Second, rare variants that introduce a gain of function will often not overlap regulatory annotations in publicly available data. An accurate model for regulatory activity can predict the gain of function, allowing follow-up consideration of the site.

In recent years, artificial neural networks with many stacked layers have achieved breakthrough advances on benchmark datasets in image analysis (Krizhevsky et al. 2012) and natural language processing (Collobert et al. 2011). Rather than choose features manually or in a preprocessing step, convolutional neural networks (CNNs) adaptively learn them from the data during training. They apply nonlinear transformations to map input data to informative high-dimensional representations that trivialize classification or regression (Bengio et al. 2013). Early applications of CNNs to DNA sequence analysis surpass more established algorithms, such as support vector machines or random forests, at predicting protein binding and accessibility from DNA sequence (Alipanahi et al. 2015; Zhou and Troyanskaya 2015). More accurate models can more precisely dissect regulatory sequences, thus improving noncoding variant interpretation. However, to fully exploit the value of these models, it is essential that they are technically and conceptually accessible to the researchers who can take advantage of their potential.

Here, we introduce Basset, an open source package to apply deep CNNs to learn functional activities of DNA sequences. We used Basset to simultaneously predict the accessibility of DNA sequences in 164 cell types mapped by DNaseI-seq by the ENCODE and Epigenomics Roadmap projects (Consortium 2012; Ernst et al. 2015). From these datasets, CNNs simultaneously learn the relevant sequence motifs and the regulatory logic with which they are combined to determine cell-specific DNA accessibility. We show that a model achieving this level of accuracy provides meaningful, nucleotide-precision measurements. Subsequently, we assign GWAS variants cell-specific scores that reflect the accessibility difference predicted by the model between the two alleles. These scores are highly predictive of the causal SNP among sets of linked variants. Importantly, Basset puts CNNs in the hands of the genome biology community, providing tools and strategies for researchers to train and analyze models on new datasets. In conjunction with genomic big data, Basset offers a promising future for human genetic research.

### Deep convolutional neural networks predict genome accessibility

DNA sequence codes for the chromatin shifts that transform cells in development and disease. We focused here on DNA accessibility due to the abundance of available data and significant association with conserved segments (Thurman et al. 2012), disease variation (Maurano et al. 2012), and eQTLs (Degner et al. 2012). The ENCODE consortium performed DNaseI-seq on 125 cell types (Thurman et al. 2012), and the Epigenomics Roadmap consortium curated an additional 39 (Ernst et al. 2015). We collected and merged these sets, resulting in 2 million sites across all cells (Figure 2A). The GENCODE v18 reference gene catalog annotates these sites as 17% promoters, 47% intragenic, and 36% intergenic. A minority 4.1-19.0% (median 8.2%) are accessible in any individual cell type, with 3.8% constitutively open in >50% of the cells. For each DNaseI hypersensitive site (DHS), we extracted 600 base pairs (bp) from the hg19 reference genome around the midpoint of each site as input to the model.

To learn the DNA sequence signals of open versus closed chromatin, we applied a deep convolutional neural network (CNN). CNNs have proven highly effective in a number of diverse tasks; this set recently includes biological sequence analysis (Alipanahi et al. 2015; Zhou and Troyanskaya 2015). As opposed to manually specifying features or performing a preprocessing step to statistically learn them, CNNs perform adaptive feature extraction to map input data to informative representations during training. The convolution operation is the engine of the CNN. In a convolution layer, the algorithm scans a set of weight matrices called filters across the input; these weight matrices learn to recognize relevant patterns (Figure 1). Prior work has demonstrated that with a sufficiently large dataset, deep neural networks can learn far more expressive and accurate models than other common approaches like random forests or kernel methods (Bengio et al. 2013).

**Figure 1.**
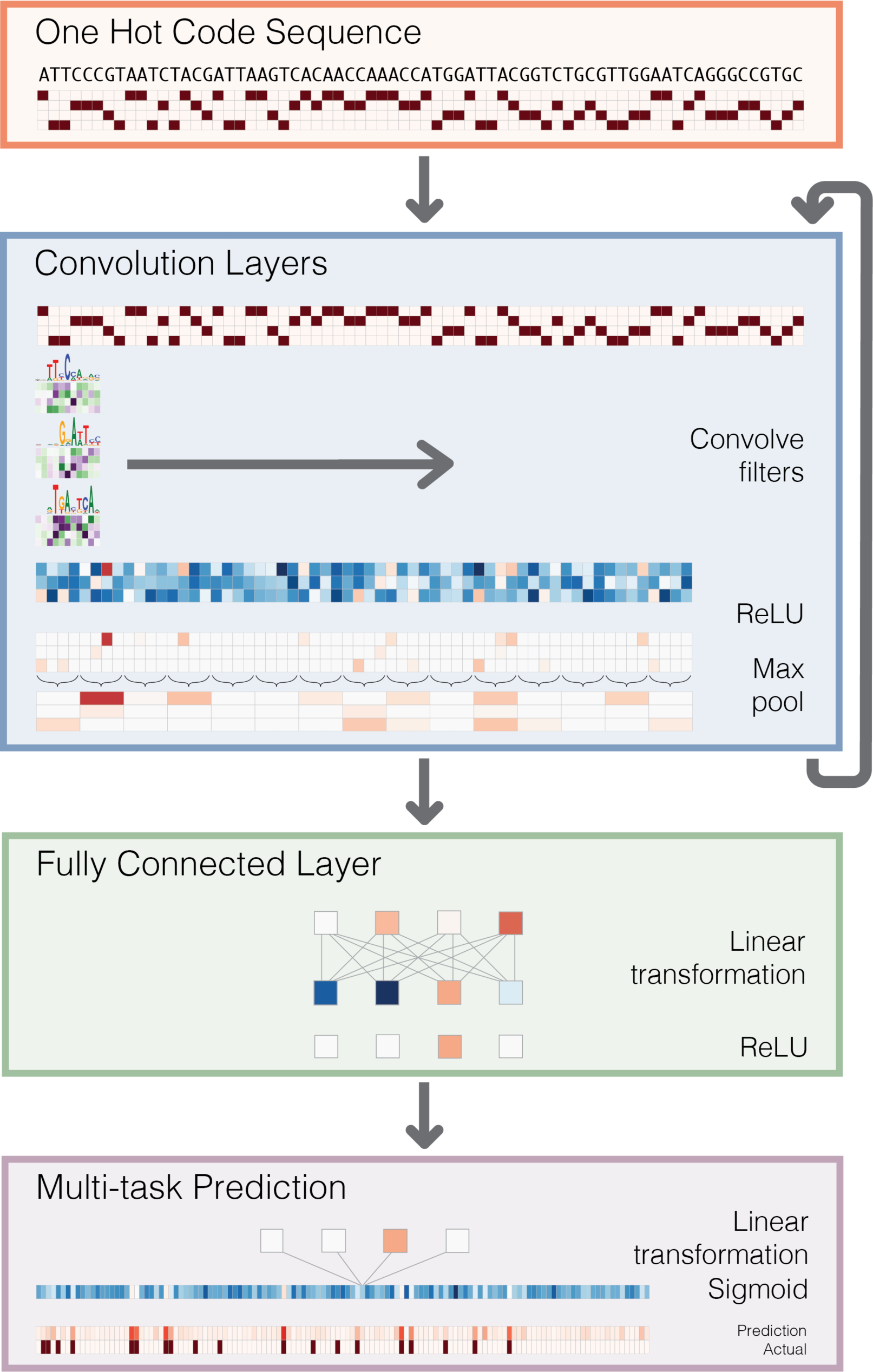
Deep convolutional neural network for DNA sequence analysis. Basset predicts the cell-specific functional activity (here DNaseI hypersensitivity) of sequences. First, we convert the sequence to a “one hot code” representation, where each position has a four element vector with one nucleotide’s bit set to one. Convolution layers proceed by scanning weight matrices across the input matrix to produce an output matrix with a row for every convolution filter and a column for every position in the input (minus the width of the filter). We apply a rectified linear unit (ReLU) nonlinear transformation to the convolution output and pool by taking the max across a window of adjacent positions. The first convolution layer operates directly on the one hot coding of the input sequence, making the convolution filters akin to the common bioinformatics tool position weight matrices. Subsequent convolution layers consider the orientations and spatial distances between patterns recognized in the previous layer. Fully connected layers perform a linear transformation of the input vector and apply a ReLU. The final layer performs a linear transformation to a vector of 164 elements that represents the target cells. A sigmoid nonlinearity maps this vector to the range 0-1, where the elements serve as probability predictions of DNaseI hypersensitivity, to be compared via a loss function to the true hypersensitivity vector.

For DNA sequences, the initial convolution layer corresponds to optimizing the weights of a set of position weight matrices (PWMs), which are a well-studied tool in bioinformatics (Stormo 2000). These PWM filters search for their motifs along the sequence and output a matrix with a row for every filter and column for every position in the sequence (Figure 1). Computing nonlinear functions of the information flowing through the network allows for more expressive models. In each convolution layer, we apply a rectifier operation (i.e. set negative values to 0) to the matrix of filter output (Nair and Hinton 2010). Finally, we pool adjacent values by taking the maximum in a small window. This operation reduces the dimension of the input to the next layer (and thus the computation required in training). It also provides invariance to small sequence shifts to the left or right.

Subsequent convolutional layers operate analogously on the output of the prior layer. Thus, for DNA sequences, they capture spatial interactions between the initial PWM filter outputs. The full architecture of our neural network includes three convolution layers and two layers of fully connected hidden nodes. In general, deeper networks are able to learn more abstract representations; here, depth allows the algorithm to consider the sequence at multiple resolutions and learn sophisticated regulatory codes that combine the recognized sequence motifs. The final layer outputs 164 predictions for the probability that the sequence is accessible in each of the 164 cell types. During training, we compare these predictions to the experimentally measured accessibilities and update the model parameters to improve the predictions (see Methods).

We have released open-source software implementing all procedures described in this manuscript, including routines to preprocess common functional genomics data, train the network, and extract the knowledge it has learned. We have named the package Basset, as an allusion to the extraordinary abilities of this hound dog to learn a scent that they are subsequently able to detect and pursue.

We trained Basset and a recently published advance based on gapped kmer support vector machines called gkm-SVM (Lee et al. 2015) to predict the accessibility of a set of test sequences in 164 cell types (see Methods). To synthesize sensitivity and specificity, which inherently trade off, we assessed the models using the area under the receiver operating characteristic curve (AUC), which plots the false positive rate versus the true positive rate. By this measure, Basset is exceptionally accurate, achieving a mean AUC of 0.895 over all cells, relative to 0.780 for gkm-SVM (Figure 2B). Basset substantially improved the AUC for every cell type. At a false positive rate of 10%, Basset identifies 55-80% of true positive DHS sequences (Figure 2C, Supplementary Figure 1). In addition to predicting constitutive sites, Basset effectively captures cell and lineage-specific accessibility (Supplementary Figure 2). Finally, Basset predicted accessibility of sites assigned to various annotation classes with consistent accuracy: 0.900 for promoters, 0.884 for intragenic, and 0.891 for intergenic (Supplementary Figure 3).

**Figure 2.**
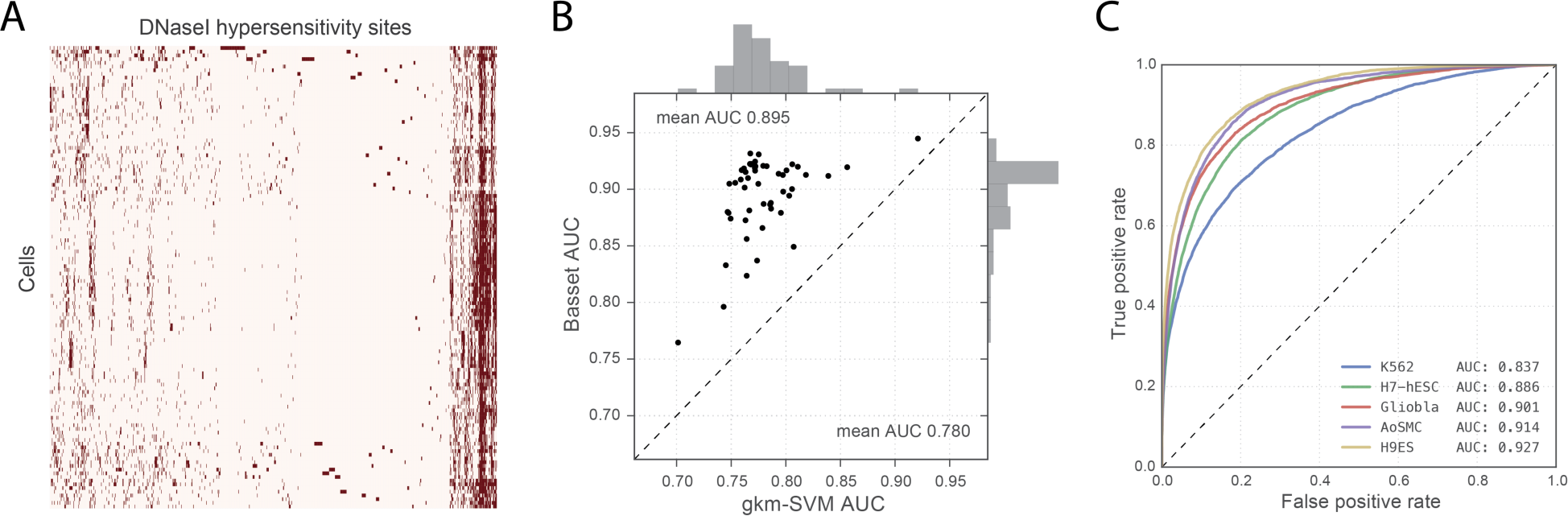
Basset accurately predicts cell-specific DNA accessibility. (A) The heat map displays hypersensitivity of 2 million DHSs mapped across 164 cell types. We performed average linkage hierarchical clustering using Euclidean distance to both cells and sites. (B) The scatter plot displays AUC for all cell types achieved by Basset and the state-of-the-art approach gkm-SVM, which uses support vector machines. (C) The ROC curves display the Basset false positive rate versus true positive rate for five cells, selected to represent the .05, .33, .50, .67, and .95 quantiles of the AUC distribution.

### Basset recovers known protein binding motifs

Though deep neural networks predict accurately, the principles that they learn are not trivially interpretable as they are in simpler linear models. However, substantial information can be extracted from the model by examining its parameters, modulating the flow of information through components of the network, and exploring its predictions on purposefully chosen sequences.

The typical DHS is a nucleosome-free region where protein(s) bind the DNA to create an accessible site (Voss and Hager 2014; Sherwood et al. 2014). Thus, we expect that a predictive model of accessibility will capture this dependence by learning the DNA binding sites of a variety of universal and cell-specific proteins. The first convolution layer of the model scans the DNA sequence with a set of pattern-recognizing filters whose weights are optimized during training. These filters are ideal for capturing this protein-binding information.

The model’s 300 convolution filters recovered an extensive repertoire of known DNA binding protein motifs (Figure 3B). To aid in their interpretation, we nullified each filter by setting its output to a constant value: its mean output over all nucleotides in the test set. This obstructs the filter from passing any information forward through the network. We considered the vector containing the change in predicted accessibility in each cell type and quantified each filter’s influence as this vector’s sum of squares. Complementing that filter-centric analysis, we performed a protein-centric analysis by introducing known protein binding motifs from the CIS-BP data database into the center of many sequences and measuring the change in predicted accessibility (Weirauch et al. 2014).

**Figure 3.**
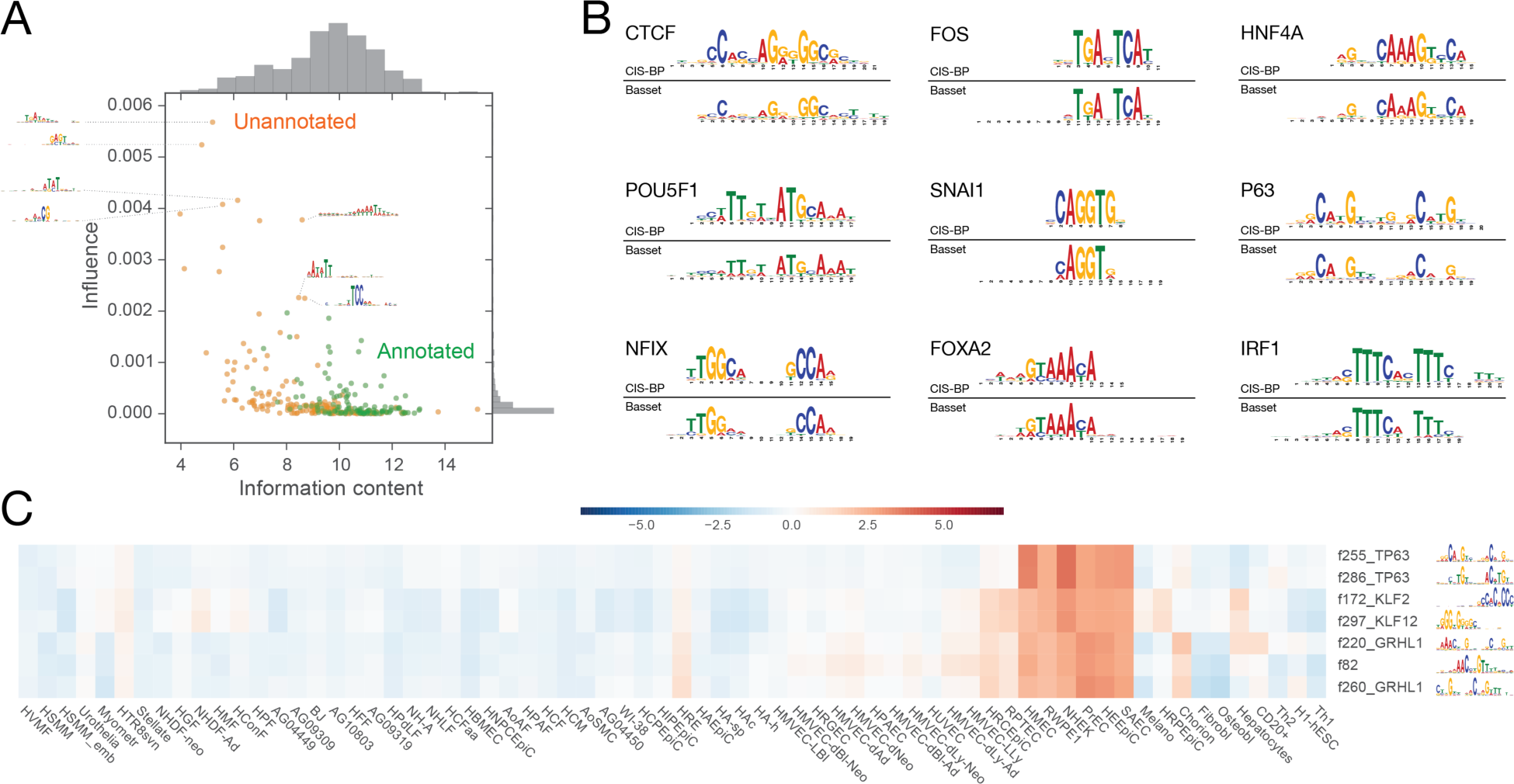
Basset initial convolutional layer discovers known and novel sequence motifs. (A) In the scatter plot, the x-axis describes the information content for the PWMs represented by the 300 first layer convolution filters. The y-axis describes an influence score, which we compute by setting all output from the filter to its mean (thus nullifying the filter) and taking the sum of squares of the vector of accessibility prediction changes over all cells. We colored filters by whether or not they could be annotated at a q-value threshold of 0.1 by the Tomtom motif comparison tool to known TF motifs in the human CIS-BP database. (B) Overall, 45% of filters could be annotated, including the alignments shown here. (C) Clustering the filters by their influence on accessibility predictions in each cell type revealed this set matching P63, GRHL1, and KLF factors, which are known to be involved in epithelial development.

The critical genome architecture protein CTCF was most predictive of accessibility across all cell types. The model dedicates the most filters (12) to comprehensively represent CTCF’s 19 bp long DNA recognition site (Supplementary Figure 4). Each filter focused on overlapping portions and variations of the motif. The AP-1 complex, consisting of proteins from the JUN, FOS, ATF, and JDP families, also emerged as highly influential via 4 filters. AP-1’s important role in regulating open chromatin has been previously observed (Biddie et al. 2011).

At a q-value threshold of 0.1, 45% of the filters aligned significantly to protein motifs in CIS-BP, which were originally acquired by independent ChIP-seq or in vitro experiments (Gupta et al. 2007; Weirauch et al. 2014). This set included highly similar weight matrices for many developmental regulators; Figure 3B depicts a sample. Many more filters capture partial coverage of known motifs, which were deemed insignificant matches to the database after multiple testing correction (Supplementary Figure 4). Motifs known to allow for variable spacing between two components were not prominently recognized by these partial motif filters (Reid et al. 2010).

Some unrecognized filters captured lower-order sequence composition, such as the known enrichment of higher GC content in TF-bound DHSs (Wang et al. 2012). One filter directly detects CpG’s, which can be methylated and are a well-studied feature of regulatory modules (Bird and Wolffe 1999). Other examples measure poly-AT stretches and non-consecutive C’s and G’s (thus avoiding the CpG). Their influence emphasizes the importance of the local sequence context of binding motifs to their function (Figure 3A) (Levo and Segal 2014; Dror et al. 2015).

To further explore how these filters influence predictions, we studied the AP-1 consensus motif TGASTCA in detail. Rather than indiscriminately predicting accessibility upon detection of the motif, Basset accurately captures the nuance of these binding events. The model achieved an average AUC of 0.86 across all cell types on test sequences that contain the motif in the center 50 nt; a classifier that had only learned the motif would merely be guessing at which specific motif instances were bound and achieve AUC 0.5.

To determine what features the model uses to vary its predictions for AP-1 motifs, we artificially inserted the motif into the center of a random set of sequences and compared predictions before and after. We focused the analysis on mammary fibroblast (HMF) and H7-hESCs, which serve as examples of cells that respond with strong and medium strength to the AP-1 motif respectively. The motif alone was insufficient to predict high accessibility, but does shift most predictions upward to varying degrees (Supplementary Figure 5). We noted several features that influence Basset’s predictions. The first nucleotide 5’ prefers [ACG] rather than T, and the first nucleotide 3’ prefers [CGT] rather than A. This is apparent in the weight matrix for the primary filter used by the model to identify AP-1 (Supplementary Figure 6). In HMFs, the average prediction for the full consensus motif is 0.49, versus 0.25 for the motif with a 5’ T, and 0.10 for the motif with a 5’ T and 3’ A.

The sequence composition of an additional +100 nt of flanking sequence also influences the predictions. The clearest effect comes from flanking poly-AT stretches that drive down the prediction (Supplementary Figure 6). So-called A-tracts are known to narrow the minor groove of the DNA double helix (Rohs et al. 2009). Additional protein binding sites also influence predictions. We observed an interesting case of a GGAART motif (best represented in CIS-BP by ETS family member FEV) that can overlap 5’ the AP-1 motif for a high prediction (Supplementary Figure 7). We also observed that nearby weak non-consensus TTASTCA AP-1 motifs increase the prediction (Supplementary Figure 7). These observations suggest that convolutional neural networks offer a simple and affective approach to automatically capture the subtle influences on functional activity provided by local sequence composition (Slattery et al. 2014).

Several unrecognized filters had high information content, which indicates that they may refer to unannotated proteins or alternative binding modes of annotated ones. To further explore these filters, we computed the influence of nullifying each filter on the downstream cell accessibility predictions (Supplementary Figure 8). Clustering these influence profiles reveals modules of filters matching proteins known to regulate development to their active cell types. For example, filters matching database motifs for known epithelial regulators P63, GRHL1, and KLF factors are predictive of accessibility in a variety of epithelial cells (Figure 3C) (Pignon et al. 2013; Wilanowski et al. 2008; Ray and Pollard 2012). The unrecognized filters span a range of cell preferences, and future work will be necessary to determine their role and potential binding proteins.

### In silico saturation mutagenesis pinpoints nucleotides driving accessibility

A trained model can be used to predict the functional activity of arbitrary sequences, offering a powerful approach to understand and apply the regulatory grammar that is has learned. Saturation mutagenesis experiments, in which every mutation to a sequence is tested, are a powerful tool for dissecting the exact nucleotides driving a functional activity (Patwardhan et al. 2009; Melnikov et al. 2012). State of the art experimental approaches involve a complex procedure of synthesizing massive pools of oligonucleotides and measuring their activity in a parallel reporter assay (Melnikov et al. 2012; Patwardhan et al. 2012). By computing the predicted accessibility of all possible mutations to a sequence, Basset can be used to perform an in silico saturation mutagenesis.

We constructed heat maps that display the change in predicted accessibility from mutation at every position to each alternative nucleotide. These maps highlight the individual nucleotides most critical to a sequence’s activity. We assigned two scores to every position: (1) the loss score measures the largest possible decrease and (2) the gain score measures the largest increase.

High loss scores mark positions with existing functional motifs where mutations can damage the motif and decrease accessibility. For example, the sequence mapped in Figure 4 is accessible in embryonic stem cells and contains the AP-1 motif. AP-1 complex members JUN and JUND ChIP-seq in H1-hESCs support the binding event, and PhyloP conservation statistics support the precise relevance of the TGASTCA motif (Pollard et al. 2010). Mutations within the motif and numerous flanking nucleotides result in decreased predicted accessibility. Genome-wide, a synthesis of loss scores from all cell types significantly correlated with PhyloP (Pearson 0.188; p-value 4.4e-102) (see Methods) (Figure 4B).

**Figure 4.**
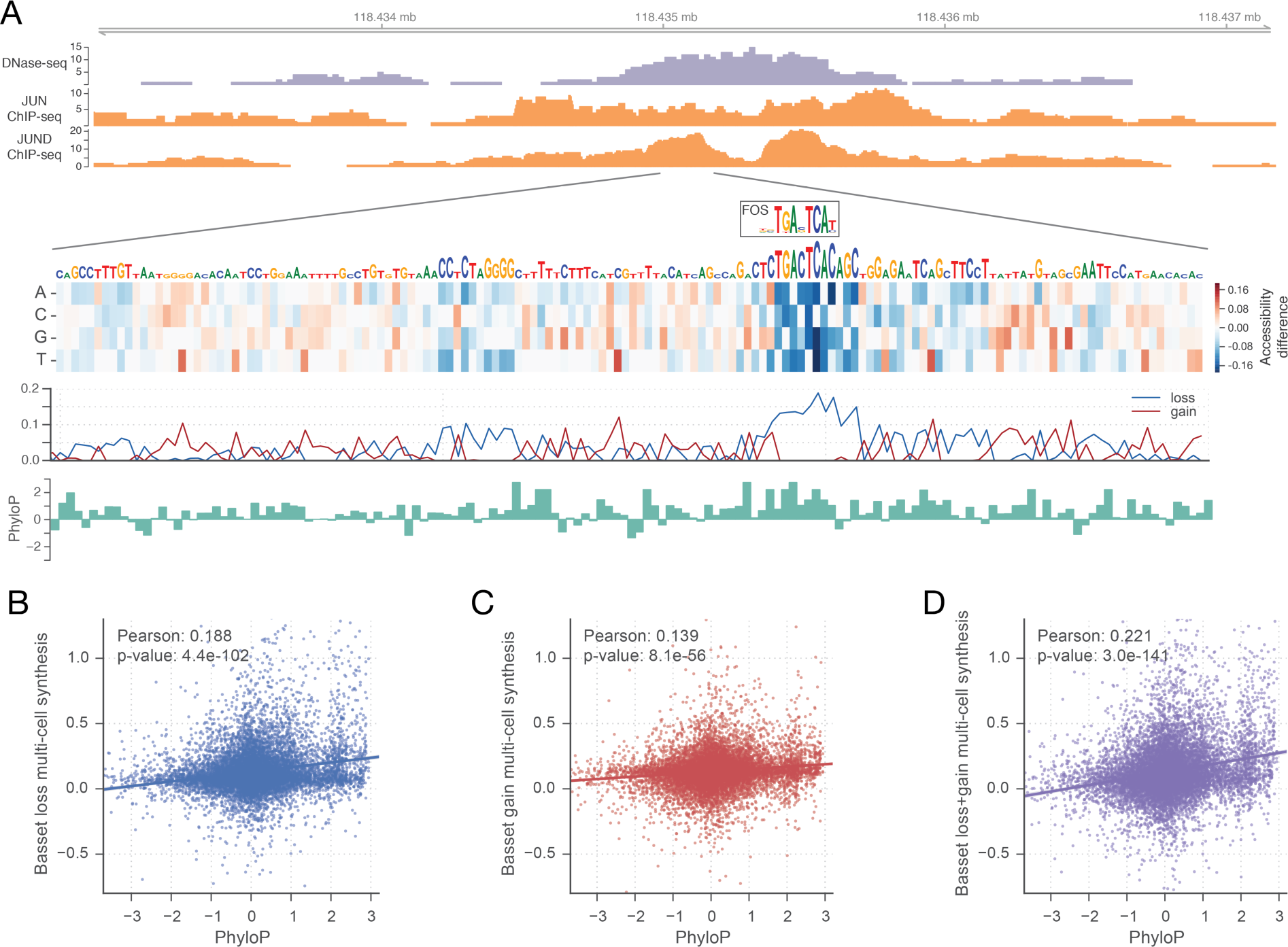
In silico saturated mutagenesis for DNaseI hypersensitivity. (A) We used Basset to predict the effect of every mutation on the accessibility of the region chr9:118,434,976-118,435,175 in H1-hESCs. The heat map displays the change in predicted accessibility for mutated sequences. Each column corresponds to a position in the sequence. Each row represents mutation to the corresponding nucleotide. In the line plot below, loss scores measure the maximum decrease among all mutations from the true nucleotide. Gain scores measure the maximum increase. We drew nucleotides to be proportional to the loss score, beyond a minimum height. At this locus, the model highlights the TGASTCA motif of the AP-1 complex (shown as the CIS-BP database motif for FOS). ChIP-seq of JUN and JUND in H1-hESCs confirm binding of the complex. The bound motif displays high conservation according to PhyloP. (B) Genome-wide, loss scores had a strong relationship with PhyloP (see Methods). (C) Gain scores alone had a weaker relationship, but the (D) combination achieved the strongest relationship.

In contrast, high gain scores suggest latent potential in a sequence; the corresponding mutation often introduces a functional motif to increase the predicted accessibility. Although such a position does not mark a present functional motif, there may be negative selection against one forming and rearranging accessibility in the region. In support of this effect, considering both loss and gain scores increased the Pearson correlation with PhyloP to 0.221 (p-value 3.0e-141) (Figure 4D). This correlation was consistent across promoter, intragenic, and intergenic annotation classes (Supplementary Figure 9).

### Basset predicts greater accessibility changes for likely causal GWAS SNPs

Genome-wide association studies (GWAS) have uncovered ample noncoding variants associated with physical traits and disease in human populations (Welter et al. 2014). DHSs are highly enriched for GWAS SNPs, which can modulate the accessibility of the site to affect local gene expression (Maurano et al. 2012; Degner et al. 2012). Basset captures the sequence signals driving accessibility and ought to have predictive power for prioritizing noncoding variants and suggesting mechanistic hypotheses for further investigation into their causal role for the phenotype. For this purpose, we defined SNP Accessibility Difference (SAD) profiles as the difference in predicted accessibility across cell types between two alleles.

The scarcity of confirmed positive examples of noncoding causal variants challenges a thorough assessment of the value of SAD scores for GWAS prioritization. Instead, we studied probabilistic assessments of causality assigned by an orthogonal method. 8741 GWAS SNPs associated with auto-immune disease were analyzed with a statistical method called PICS that leverages dense genotyping data to assign a probability of being the causal SNP among a nearby set of SNPs in linkage disequilibrium (LD) (Farh et al. 2015). In a large number of cases, PICS identified the causal SNP with high probability.

We focused on a set of 7252 GWAS SNPs for which no SNP in LD affects a protein coding gene and classified 235 high-PICS SNPs that were assigned causal probability >=0.5 and 3004 low-PICS SNPs that were assigned causal probability <=0.05. SAD profile means were significantly greater for the set of high-PICS SNPs (Mann-Whitney U Test, p-value < 1.3e-7) (Supplementary Figure 10). >7 times more high-PICS SNPs than low were predicted to change accessibility by an average of >0.1 over the cell types (Figure 5A). Coverage of the SNPs in this set is wide—31% of all index SNPs had at least one SNP in its LD set for which the model predicted a >10% change in probability of accessibility in some cell type (Supplementary Figure 10). We report all predicted mutation effects on this dataset in Supplementary Table 1.

**Figure 5.**
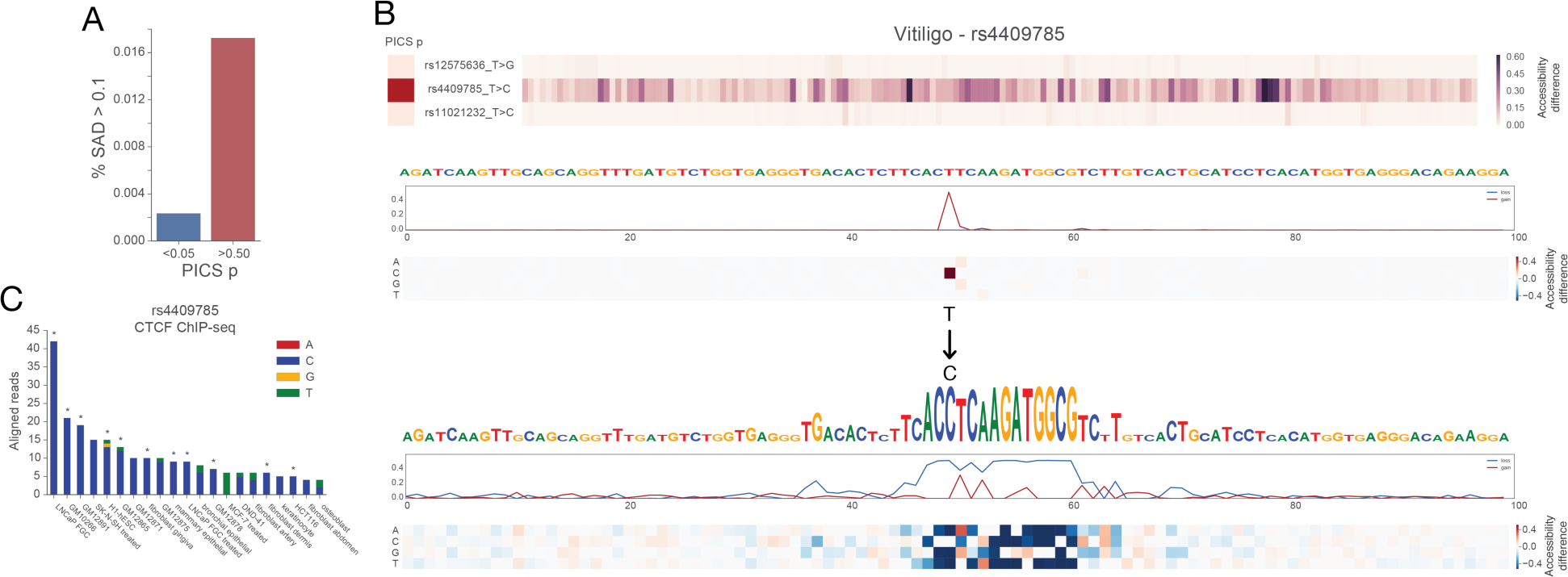
SNP accessibility difference scores enable genomic variant interpretation. (A) Basset assigned greater scores to likely causal GWAS SNPs (PICS probability > 0.5) versus unlikely nearby SNPs (PICS probability <0.05) as determined by population fine mapping data. The bars measure the proportion of SNPs assigned a SAD profile mean across all cell types >0.1. (B) We annotated rs4409785 among the highest SAD scores, in agreement with the PICS view of this haplotype block. Basset predicts the more common T allele to be completely dormant, but the region transforms with the C allele into a site deemed by Basset to have very high accessibility due to a CTCF binding site. (C) CTCF ChIP-seq in 88 unique cell types strongly supports the allele-specificity of CTCF at this site. We plotted cells with >3 reads (summed across replicates) aligned to the site, and marked significant peak calls with *’s. The 11 cells with significant peak calls all sequenced the C allele.

Among the agreements with PICS was rs4409785, associated with vitiligo (Jin et al. 2012), rheumatoid arthritis (Okada et al. 2013), and immune mechanisms in multiple sclerosis (Sawcer et al. 2011). PICS assigned rs4409785 85.3% probability of causality for vitiligo. The SNP is located in a 559 Kb gene desert. However, it has been hypothesized to regulate *TYR*, which although 6.28 Mb away, offers a plausible mechanism for the skin color disease vitiligo. *TYR* catalyzes conversion of tyrosine to melanin, the pigment that gives skin its color (Jin et al. 2012).

Basset predictions support this hypothesis; the more prevalent T allele is devoid of activity, but the C allele creates a motif recognized by the model’s CTCF filters (Figure 5B). Although this sequence imperfectly matches the CTCF database motif, Basset predicts dramatically increased accessibility in all cell types, including an increase in H1-hESCs from 0.8% probability to 73.24%. To assess experimental evidence for allele-specific CTCF binding, we downloaded 88 unique CTCF ChIP-seq datasets performed in a variety of cell types by ENCODE (Consortium 2012). Of these, 21 cell types had >3 reads aligned to rs4409785, of which 11 had significant peak calls (Figure 5C). Every significant peak sequenced the C allele, strongly supporting the allele specificity of CTCF binding at this site.

Recent work has demonstrated that modulating CTCF binding at topologically associating domain boundaries can dramatically affect gene expression by altering the architecture of the genomic region (Lupiáñez et al. 2015; Guo et al. 2015). By modeling the underlying sequence driving the experimental measurement, Basset’s highlight of rs4409785 suggests a functional hypothesis to accelerate study of this widely influential variant.

### Leveraging large-scale models allows accurate and efficient prediction of new datasets

To make the most of machine learning to identify causal SNPs, researchers must train models on data from the appropriate (perhaps specialized) cell types (Lee et al. 2015). Such cells will often be overlooked by large-scale mapping projects, but experimentally characterized by individual labs. We designed Basset to accommodate this frequent and critical scenario. We also hypothesized that Basset can rapidly learn to accurately predict new data by leveraging a model pre-trained on public data.

To test this hypothesis, we removed 15 datasets spanning the range of AUC from the pool of 164 studied above. We re-trained on the remaining 149 “public” datasets to establish a “pre-trained” model. This model achieved an average AUC of 0.892 across the 149 cells, on par with the full model analyzed above.

For each remaining dataset individually, we sampled a matching number of sites from the “public” 149 to serve as negative examples. Rather than train from scratch on the dataset alone, we initialized the model parameters with those from the “pre-trained” model. By providing this head start, we need only perform one training pass through the new data to achieve models with predictive accuracies rivaling those from the full 164 cell model above (Figure 6; Supplementary Table 2). We could complete this training procedure in an average of 6.5 hours on a Macbook 2.8 GHz Intel core i7 or 18 minutes on an NVIDIA Tesla K20m GPGPU. Thus, this approach allows researchers to train highly accurate deep CNN models on common computer hardware in a few hours.

**Figure 6.**
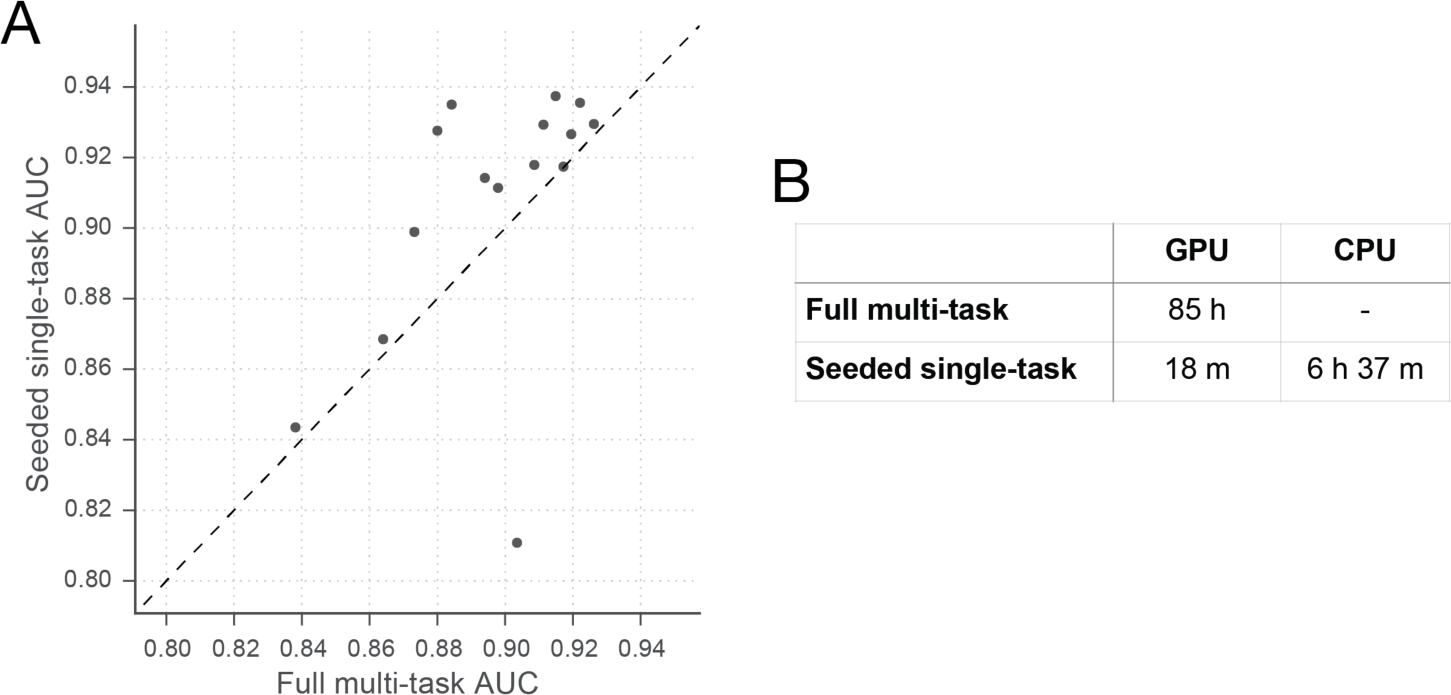
Basset leverages large-scale public data to inform additional dataset learning. The scatter plot shows AUC for 15 datasets achieved by the full model trained on all 164 cell types on the x-axis and AUC achieved by a procedure to simulate studying that dataset alone on the y-axis. To study the dataset alone, we pre-train a model on 149 cells (after removing these 15), seed training of the additional cell with that model’s parameters, and perform a single training pass through the new data. This rapid procedure was effective for all but one dataset (HRCEpiC, renal cortical epithelial cells), for which multi-task training with the many other similar epithelial cells was beneficial. The AUC improvement for many cells suggests that our full model may benefit from increased capacity or decreased regularization.

### Discussion

In this work, we introduced Basset—an open source package to apply deep CNNs to learn DNA sequence activity. Basset effectively learned the complex code of DNA accessibility across many cell types and substantially surpassed the predictive accuracy of the present state-of-the-art. We demonstrated how our model precisely implicates the nucleotides driving activity, highlighting genomic positions with either fragile activity that can be lost by mutation or latent potential activity that can be unlocked by mutation. These sites are more conserved and their mutation is more likely to cause disease.

Predicting the functional output of DNA sequences is a fundamental problem in computational biology with a long history of approaches (Bussemaker et al. 2001; Beer and Tavazoie 2004; Segal et al. 2003). Our work, considered with independent successes in other groups (Alipanahi et al. 2015; Zhou and Troyanskaya 2015), indicates that hardware (GPGPUs), software (convolutional neural networks), and training data (mammalian genome-wide) have now converged to enable drastically enhanced performance on such problems. What has been missing from our previous models? One primary difference is that convolutional neural networks naturally consider positional relationships between sequence signals. In contrast, most sequence kernel approaches immediately throw away position information to represent DNA as vectors of k-mer counts (Ghandi et al. 2014). Position-specific sequence kernels exist, but they greatly increase the dimensionality of the raw input space to which the sequence is initially mapped. Unfortunately, such kernel methods do not scale well to higher dimensional inputs, both in computational efficiency and predictive performance, due to the curse of dimensionality—generalization via interpolation suffers in higher dimensional spaces. By jointly learning a meaningful representation and a smooth parameterized projection to the outputs, convolutional neural networks, in essence, *learn the kernel*. Finally, neural networks trained via stochastic gradient descent scale very well to large datasets, allowing us to learn good parameters within a general and expressive model structure.

The most successful prior approaches to analyze noncoding variants compare them to the broad regions that functional genomics experiments have annotated to have reproducible accessibility, protein binding, and/or histone modifications (Kircher et al. 2014; Fu et al. 2014; Ritchie et al. 2014). Basset has two primary benefits over this approach. By directly modeling the mapping from sequence to activity, Basset implicates the precise nucleotides influencing accessibility, providing a finer resolution view than mere overlap with a broad region. Basset assigns low SAD profiles to many nucleotides overlapped by these regions, calling into question their consideration for causal roles.

Furthermore, if the genome sequenced in the original experiment included only the inaccessible allele, there will be no indication that accessibility is relevant to the SNP. Basset readily identifies these gain-of-function mutations, as demonstrated for CTCF binding to rs4409785 (Figure 5). As statistical searches for influential variants in human populations continue, rare variants will make up a greater proportion. Especially for these rare variants, the functional genomics experiment will be unlikely to have been performed in the necessary genetic background. This growing trend makes methods like Basset even more important if we are to interpret these variants.

With Basset, a researcher can perform a single sequencing assay in their cell type of interest and simultaneously learn that cell’s chromatin accessibility code and annotate every mutation in the genome with its influence on present accessibility and latent potential for accessibility. By leveraging large-scale public data, one can train highly accurate models on common computational hardware. Researchers continue to discover noncoding variants in human populations that influence phenotypes, and such annotation will be indispensable for interpreting how those variants function. As the tide of functional genomics data continues to flow, novel machine learning approaches such as deep convolutional neural networks have great power to aid this goal.

## Methods

### DNasel hypersensitivity data

We downloaded DNaseI-seq peak BED format files for 125 cell types from the ENCODE project and 39 cell types from the Epigenomics Roadmap project. For both sets, the previous groups called peaks using the HotSpot algorithm and performed a simulation procedure to establish a set with 1% false discovery rate. We considered all peaks, regardless of overlap with genomic annotations.

To merge the peaks into one set, we first extended each one from its midpoint to 600 bp. We greedily merged peaks based on their distance to an adjacent peak until no peaks overlapped by more than 200 bp. During a merger of peaks, we specified the activity of the new peak as the union of the sets of active cell types for each individual peak. We specified the new limits by extending from a weighted average of the two peak midpoints, weighted by the number of cells each individual peak was active in. This produced a set of 2,071,886 peaks of which 4.1-19.0% (median 8.2%) were active in the individual cell types. We extracted the hg19 reference genome sequence for each merged site as input to the model.

For some analyses of model predictions, we divided sites between promoter (within 2 kb of a transcription start site), intragenic (overlapping a gene’s span), and intergenic classes using the GENCODE v18 reference catalog (Harrow et al. 2012).

### Deep convolutional neural network

Deep convolutional neural networks (CNNs) are a type of deep neural network that are specifically parameterized to take advantage of known spatial structure. They were originally developed to recognize handwritten digits in images (LeCun et al. 1998). Convolutional networks have since become the gold standard for numerous image analysis tasks (Krizhevsky et al. 2012; Szegedy et al. 2014). Recently, convolutional networks have been modified for use within natural language processing and text analysis by applying a one dimensional convolution temporally over a sequence (Hu et al. 2014; Zhang and LeCun 2015).

We implemented a deep CNN using Torch7 (http://torch.ch). Initially, we map the DNA sequence to 4 rows of binary variables representing the presence or absence of an A, C, G, or T at each nucleotide position. The first convolutional layer of the network scans position weight matrices across the sequence (Figure 1). The matrix weights are parameters learned from the data. These are typically referred to as filters in the CNN literature. After convolving the matrix across the sequence, we applied a rectified linear ReLU nonlinearity (f(x) = max(0,x)), which has been found helpful in avoiding the vanishing gradient problem that plagued early deep learning research (LeCun et al. 1998; Nair and Hinton 2010). Finally, we “pool” adjacent positions by taking the max from a small window in order to reduce the number of parameters and achieve invariance to small shifts of the sequence left or right.

Subsequent convolutional layers operate on the output of the prior layer, which represents recognition of filter patterns across windows of the sequence. After three convolutional layers, we placed two standard fully connected artificial neural network hidden layers and a final fully connected sigmoid transformation to 164 outputs, representing the predicted probability of accessibility in each cell type. We trained to minimize the binary cross entropy loss function, summed over these 164 outputs.

We applied stochastic gradient descent to learn all model parameter, including those representing convolution filters, using RMSprop updates on mini-batches (Tieleman and Hinton 2012). First, we randomly initialized the parameters to small values. During training, the network computes predictions for small batches of sequences. We compare these predictions to the true experimental measurements using the loss function. We then update the model parameters to improve those predictions by taking a step in the direction of the gradient of the parameters with respect to the loss function, which we compute using the back propagation algorithm. After iterating over many batches of training data, the model begins to recognize specific sequence motifs indicative of accessibility and project this recognition through the network to the cell predictions. We continue training until accuracy ceases to increase on a held out validation set for 12 passes through the training data.

The user must specify the number of each type of layer, number of filters per convolution layer, filter sizes, pooling widths, fully connected layer units, and numerous regularization and training optimization parameters. We experimented with various model architectures and hyper-parameter settings using Bayesian optimization, implemented in the package Spearmint (available from https://github.com/HIPS/Spearmint) (Snoek et al. 2012). We committed to analyzing a top performing architecture that is depicted in Supplementary Figure 11. Importantly, we apply batch normalization after every layer, which substantially stabilized training optimization (Ioffe and Szegedy 2015).

Source code implementing all steps—date preprocessing, training, and downstream analysis—is available in the package Basset from http://www.github.com/davek44/Basset.

### Training, validation, and test datasets

From the 2,071,886 total sites, we reserved 71,886 for testing and 70,000 for validation, leaving 1,930,000 for training. We performed optimization directly on the training set. We used the validation set for “early stopping” after 12 epochs of unimproved validation loss and Bayesian optimization. We performed all assessment and analysis on the test set.

### gkm-SVM

We downloaded gkm-SVM v1.3 from *http://www.beerlab.org/gkmsvm/* (Lee et al. 2015; Ghandi et al. 2014). Because the code computes the full Gram matrix, we could only feasibly train on a 100k subsample of the full dataset. For each cell type, we downsampled the inactive sequences to match the number of active sequences. Though Basset easily handles imbalanced datasets, we found the natural imbalances of this DHS dataset significantly decreased gkm-SVM accuracy. Using default options, gkm-SVM required 16 days to train and test on 50 randomly selected cell types.

### Motif comparison

We identified the likely binding protein for Basset-learned first convolution layer filters by querying the CIS-BP motif database (Weirauch et al. 2014) (accessed on June 12, 2015) using the TomTom v4.10.1 search tool (Gupta et al. 2007) and requiring an FDR q-value < 0.1.

### Comparison of Basset predictions to PhyloP

We used Basset to compute loss and gain scores for every nucleotide. We compute the loss score as the predicted activity with the reference nucleotide subtracted by the minimum predicted activity after mutating the position to the alternative three nucleotides. We compute the gain score as the maximum predicted activity after mutation subtracted by the reference nucleotide activity. To ask whether these scores have a significant statistical relationship with nucleotide conservation, we required a method to consider all 328 scores per nucleotide with respect to PhyloP. We applied a linear regression model with ridge penalty, training on 80% of the nucleotides and testing on the remaining 20%. We limited the analysis to nonrepetitive regions where more confident PhyloP statistics can be assigned and to the center 100 bp of the DHSs, where Basset makes stronger predictions (Supplementary Figure 12).

### PICS

We downloaded 8741 PICS SNP annotations from the supplement of the authors’ manuscript (Farh et al. 2015). For SAD profile comparison, we focused on an unquestionably noncoding set of 7252 SNPS by removing all SNPs linked to a SNP in a protein coding gene. Without sufficient training data to learn weights for the various cell types studied, we resorted to comparing the SAD profile means to the PICS causal SNP probabilities.

## Competing interest

The authors declare that they have no competing interests.

## Acknowledgments

The authors acknowledge Abbie Groff, David Hendrickson, Marta Mele, Stephanie Fine Sasse, Chinmay Shukla, and Michael Ziller for feedback on the manuscript. DK was supported by NIH K25 award ES022984-02.

